# The PDF Data Extractor (PDE) Pre-screening Tool Reduced the Manual Review Burden for Systematic Literature Reviews by Over 35% Through Automated High-Throughput Assessment of Full-Text Articles

**DOI:** 10.1101/2021.07.13.452159

**Authors:** Erik Stricker, Michael E. Scheurer

**Author notes:** **For correspondence:** (ES); (MES). Department of Pediatrics, Department of Molecular Virology and Microbiology, Baylor College of Medicine, Houston, TX 77030, USA.

## Abstract

Literature reviews are generally time-consuming and rely heavily on accurate representation of the data in the title and abstract of articles. Often minor results and details are lost in a systematic screen, which is becoming even more frequent with the rapidly rising numbers of daily published scientific articles. We developed the PDF Data Extractor (PDE) R package to aid scientists at any stage in literature reviews while offering a user-friendly interface. The tool permits the user to categorize large numbers of full-text articles in PDF format, export containing tables to Excel sheets (pdf2table), and extract relevant data using a simple user interface, requiring no bioinformatics skills. Specific features of the literature analysis comprise the adaptability of analysis parameters including the use of regular expressions, machine learning-powered detection of abbreviations of search words in articles, and the export of document meta-data. We exemplify how the PDE R package can be utilized as a pre-screening tool allowing automated categorization of full-text articles by relevance, thereby reducing the literature to be evaluated (in our example by 35% with a sensitivity of 100% at standard parameters). The PDE R package is available from the Comprehensive R Archive Network at https://CRAN.R-project.org/package=PDE and as web tool with limited capacity at https://erikstricker.shinyapps.io/PDE_analyzer/.

## Introduction

In an age of exponentially increasing numbers of published scientific articles, it is surprising that most systematic literature reviews are still conducted manually evaluating each article individually. Systematic literature reviews aim to find and collect relevant information concerning a specific research question and are an essential step in virtually every area of research, e.g., for the preparation of review articles, project proposals, and experimental designs. While machine learning tools are available for literature searches and screens (***Marshall and Wallace, 2019***), they: 1) require a large number of manually evaluated articles for the training of the tool, 2) are often restricted to filtering articles by study design or choosing topics from a limited set of terms, and 3)are generally limited to the evaluation of article titles and abstracts.

Furthermore, extraction of tabularized data from PDF files poses an additional challenge. Existing R packages such as pdftools (***Ooms, 2019***) or tabulizer (***Leeper, 2018***) require programming and data conversion skills, deterring many users. In addition, table extraction with these packages requires the tables to either be the only element on a page or be manually selected for each instance. While many online tools allow the fast and relatively accurate conversion of PDF files into Microsoft Excel files, they require a paid subscription for the processing of more than a few files (e.g., https://smallpdf.com/, https://www.adobe.com/acrobat/online/pdf-to-excel.html, https://pdftables.com/) or are limited in input file numbers and, therefore, do not allow high-throughput table extraction (e.g., https://pdftoxls.com/, https://docs.zone/pdf-to-excel, https://www.pdftoexcel.com/).

Addressing these limitations, we developed the openly available, free-to-use PDE R package with user-friendly interface. The PDF data extractor (PDE) R package easily extracts information and tables from full-text articles in Portable Document Format (PDF) format based on user-defined keywords and does not require a training set. The output tables are saved in universal comma-separated values (.csv) or tab-separated values (.tsv) files easily opened in Microsoft Excel. Additional features include the adaptability of analysis parameters including the use of regular expressions, machine learning powered detection of abbreviations of search words in articles, and the export of document meta-data. The PDE R package is available from the Comprehensive R Archive Network at https://CRAN.R-project.org/package=PDE).

## Results

### Implementation of PDE R Package as Pre-screening Tool Significantly Reduced Manual Screening Requirements

In their Cochrane systematic review, Kraal *et al.* searched the Cochrane Central Register of Controlled Trials (CENTRAL; the Cochrane Library 2016, Issue 3), PubMed (1945 to 25 April 2016) and Embase (Ovid) (1980 to 25 April 2016) for articles on 131I-MIBG and HR NBL (***Kraal et al., 2015***). All article titles and abstracts were manually evaluated by the group and eligible articles were assessed in the full text. To simplify comparisons between the traditional and a PDE-facilitated approach, we concentrated our evaluations on the articles found in the PubMed database which were 2,291 out of the 3,366 articles screened by Kraal *et al.* (see Fig. 1). We were able to obtain 1262 full-text articles in PDF format and used them for pre-screening facilitated by the PDE R package. The analysis took approximately 1.5 hours with an average of 4 seconds per article and yielded 785 articles containing the filter/search words detected at least 20-times and 24 articles without containing text that could be processed, i.e., secured articles or scans of full articles. Accordingly, over one third of the articles (35.9%, 454/1262) were excluded without the requirement of a manual review. The 762 articles not excluded encompassed 38/39 (97.4%) of the PubMed-derived articles that ***Kraal et al. (2017***) assessed for eligibility. The 39 evaluated articles contained on average, 88 detected instances of filter words, with a range of 11 to 253, and an average of 56 sentences with search words extracted, ranging from 14 to 135. The two articles by ***de Kraker et al. (2008*)**, and ***Kraal et al. (2015*)**, finally included in the Cochrane systematic review displayed the filter words 72 and 89 times, with 50 and 31 sentences extracted, respectively.

**Figure 1.**
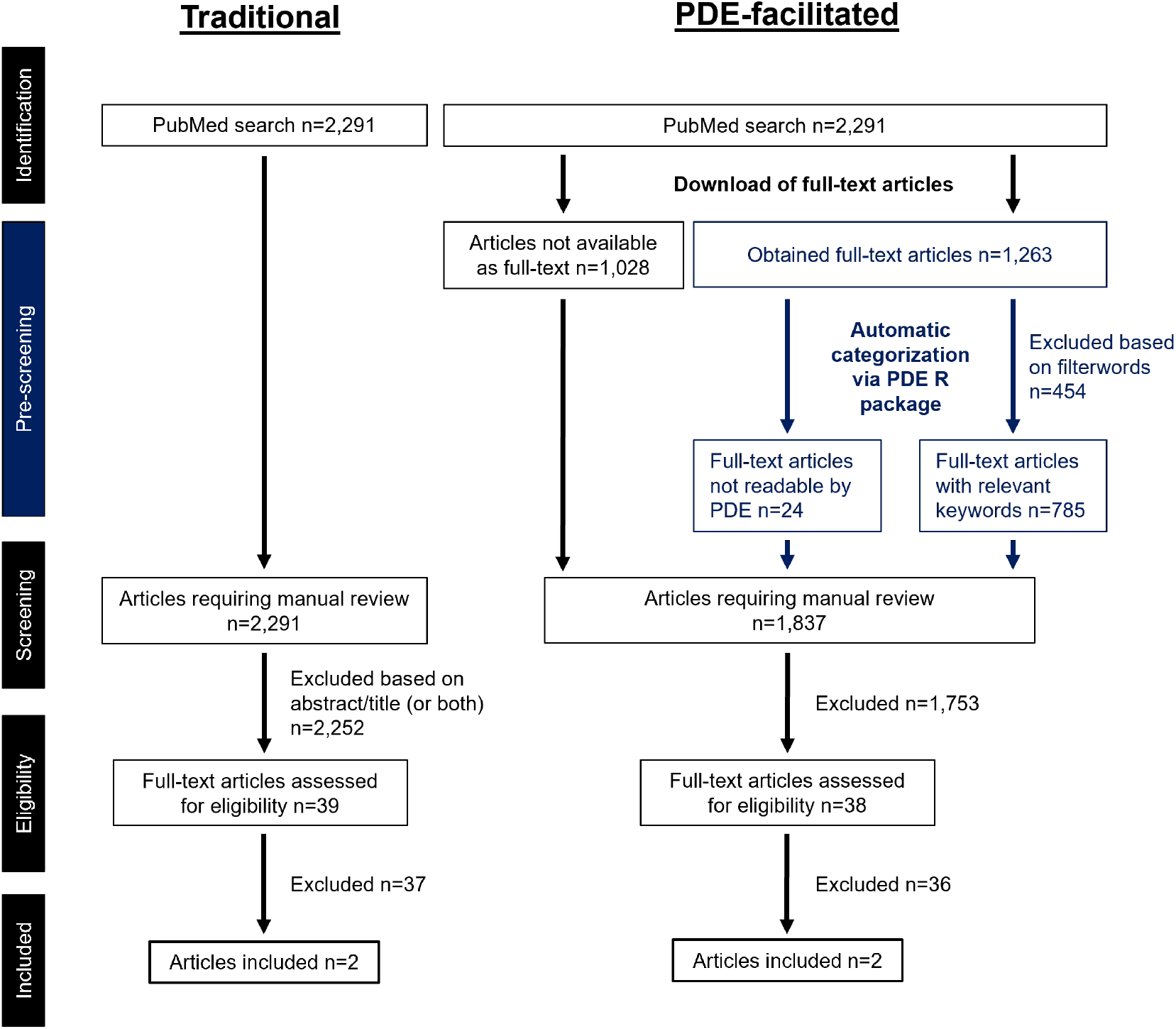
Modified PRISMA diagram comparison between the traditional approach used by ***Kraal et al. (2017*)** and the methodology using the PDE R package as prescreening tool (blue).

### The PDE R Package Displayed High Sensitivity in the Categorization of Full-Text Articles

An evaluation of specificity and sensitivity of the PDE R package was challenging since we observed that with higher thresholds for filter words, the selectivity was increased, whereas the number of articles assessed for eligibility in this pool were reduced although both included articles were still detected (Fig. 2). For example, a filter word threshold of 50 resulted in 376 articles for manual screening with and overlap of 19/39 full-text articles assessed by ***Kraal et al.(2017*)** and both included articles still being detected. Overall, at standard parameters, we can report a specificity of 64% and sensitivity of 97-100% depending on whether full-text articles assessed for eligibility by ***Kraal et al. (2017*)** or articles included in the review are considered true positives. In summary, the PDE R package notably reduced the number of articles required to be manually reviewed with high accuracy, provided additional preprocessed content for the assessment of articles, i.e., extracted sentences with search words, and evaluated articles in a reproducible manner easily applied to additional literature.

**Figure 2.**
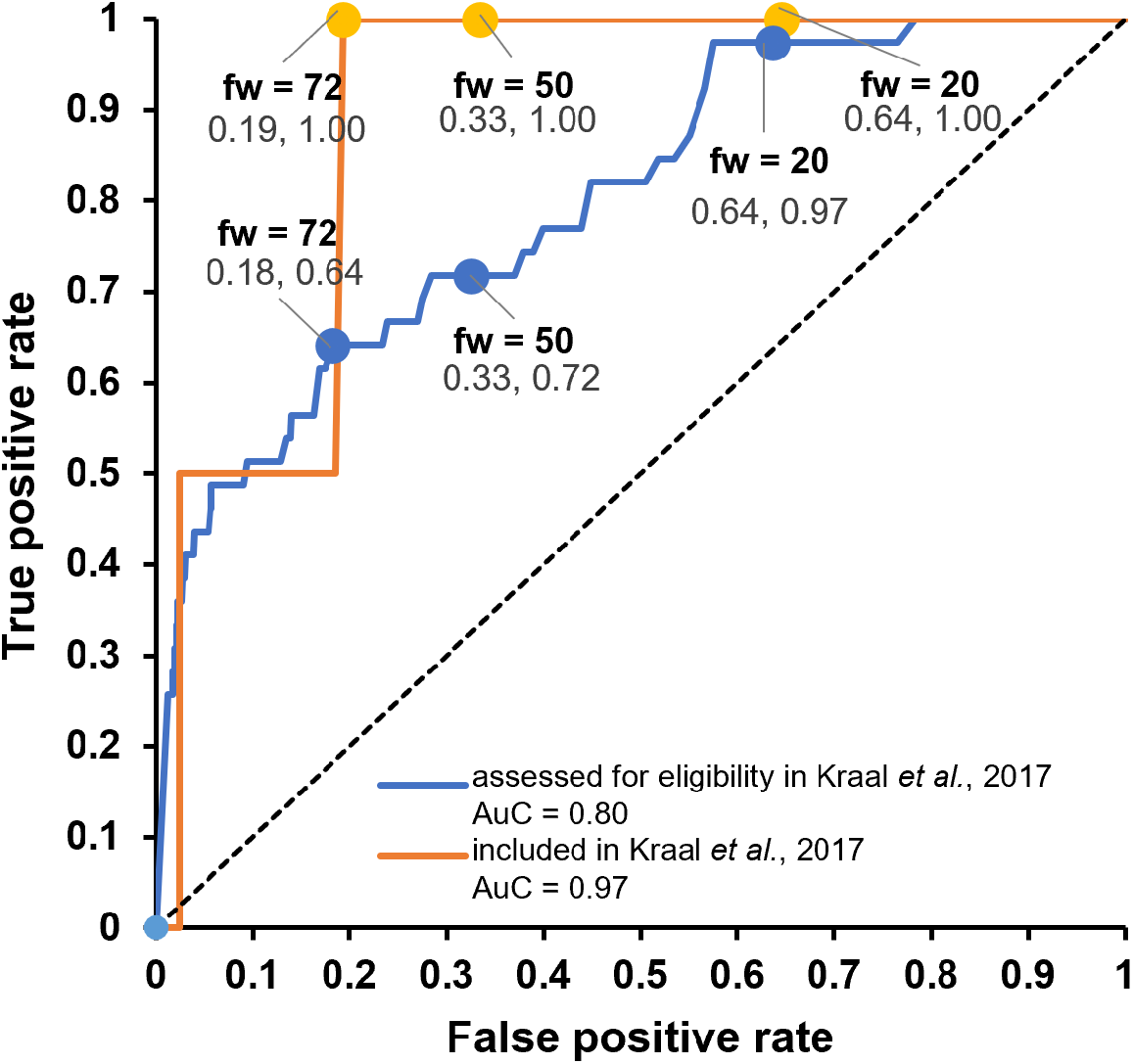
Receiver operating characteristic (ROC) curves of the article detection by the PDE_analyzer. ROC curves for the detection of the articles assessed for eligibility in ***Kraal et al. (2017*)** (blue) and the articles included in the review by ***Kraal et al. (2017*)** (orange) were compared to the reference line (dashed line). Variable parameter was the filter word threshold (fw) with different thresholds indicated by solid circles. The area under the curve (AuC) for each evaluation is indicated in the legend.

Interestingly, the article by ***Mastrangelo (1987*)** was further evaluated by ***Kraal et al. (2017*)** but then excluded from the review due to a lack of primary data. The PDE R package, on the other hand, excluded the article during initial analysis based on the number of filter word number (fw=11 was below the threshold of 20) (see Supplemental Table S1). In contrast, the clinical study by ***Garaventa et al. (1999*)** was excluded from the Cochrane Review even though the abstract and title indicated 30 patients with stage 4 neuroblastoma while the PDE R package detected 127 filter words present. Furthermore, review of the 63 sentences extracted by the PDE R package highlighted observed adverse effects such as acute myeloid leukemia, thrombocytopenia, severe interstitial pneumonia, and reverse thyroid reserve (Supplemental Table S2) (***Garaventa et al., 1999*)**. Only when consulting the clinical table in ***Garaventa et al. (1999*)** it becomes evident that only four patients with stage 4 neuroblastoma displayed MYCN amplification with >8 copies thus not reaching the inclusion criteria of at least 10 patients with HR NBL in the study. Accordingly, it cannot be excluded that other studies were falsely excluded while the PDE R package would have prevented such a mistake.

Direct comparison of abstract- and title-only review and the PDE R package showed that 97.2% (70/72) of the search words were located within the body article, only two search words in the abstract and none in the title (Fig. 3). The distribution of search words was even throughout the article by (***Mastrangelo, 1987*)** with 4/5 figures/tables including search words. While 59 search words were located in the body of the article, 11 search words were in the references. The search term “surviv” was most common with 10 incidences (8 survival, 1 surviving, 1 survivors) although absent in title and abstract.

**Figure 3.**
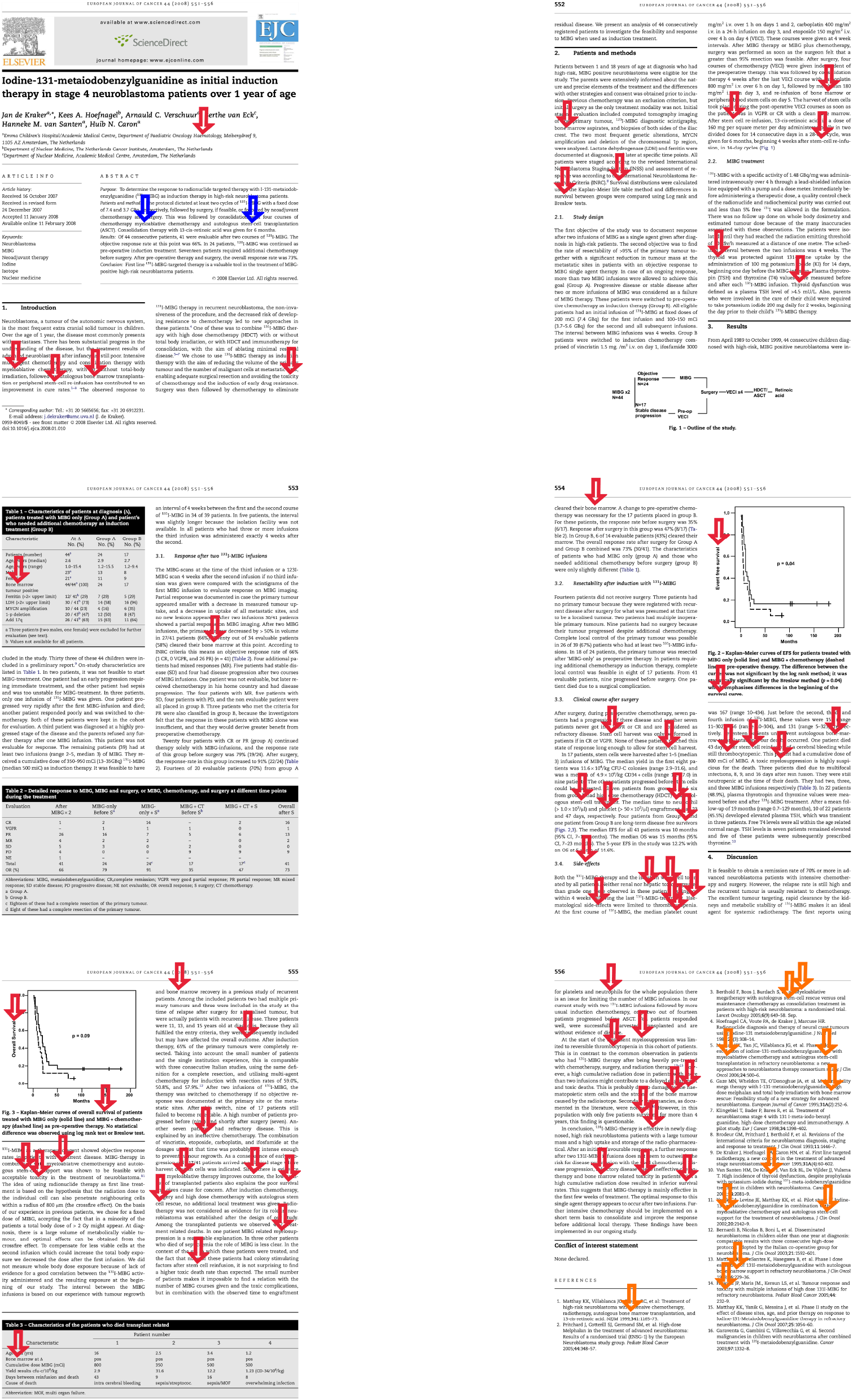
Overview of the full-text article by ***de Kraker et al. (2008*)** with highlighted search words. Blue arrows indicate the 2 search words identified in the abstract;red arrows point to the 59 search words found in the body of the article;orange arrows highlight the 11 search words detected in the references. For the list of search words see Supplemental Table S3.

### The PDE R Package Allows High-Throughput Table Extraction from PDF Files

Tables of a PDF document can be exported into a Microsoft Excel readable file format with and without the use of search words as exemplified in Figure 4. The PDE_analyzer detects the beginning of a table based on the standard annotation of tables in scientific literature, i.e., “Table” [Table index] [Table heading], or user-defined table headings, allowing the exclusion of non-table content from the PDF file. Tables that carry their heading below the actual table are not detected correctly due to the nature of the detection algorithm. To detect the content of each table cell and its location, an HTML copy of the PDF file is generated of which x and y coordinates for each text element can be extracted. For the assessment of the table structure, content starting with the fifth element in the table are compared in their coordinates. For this reason, tables with less than 5 elements cannot be processed by the PDE R package. After extraction of the table itself, the table legend is identified using font size differences in the document. The final tables are saved either as a comma- or tab-separated values file, i.e., *.csv or *.tsv.

**Figure 4.**
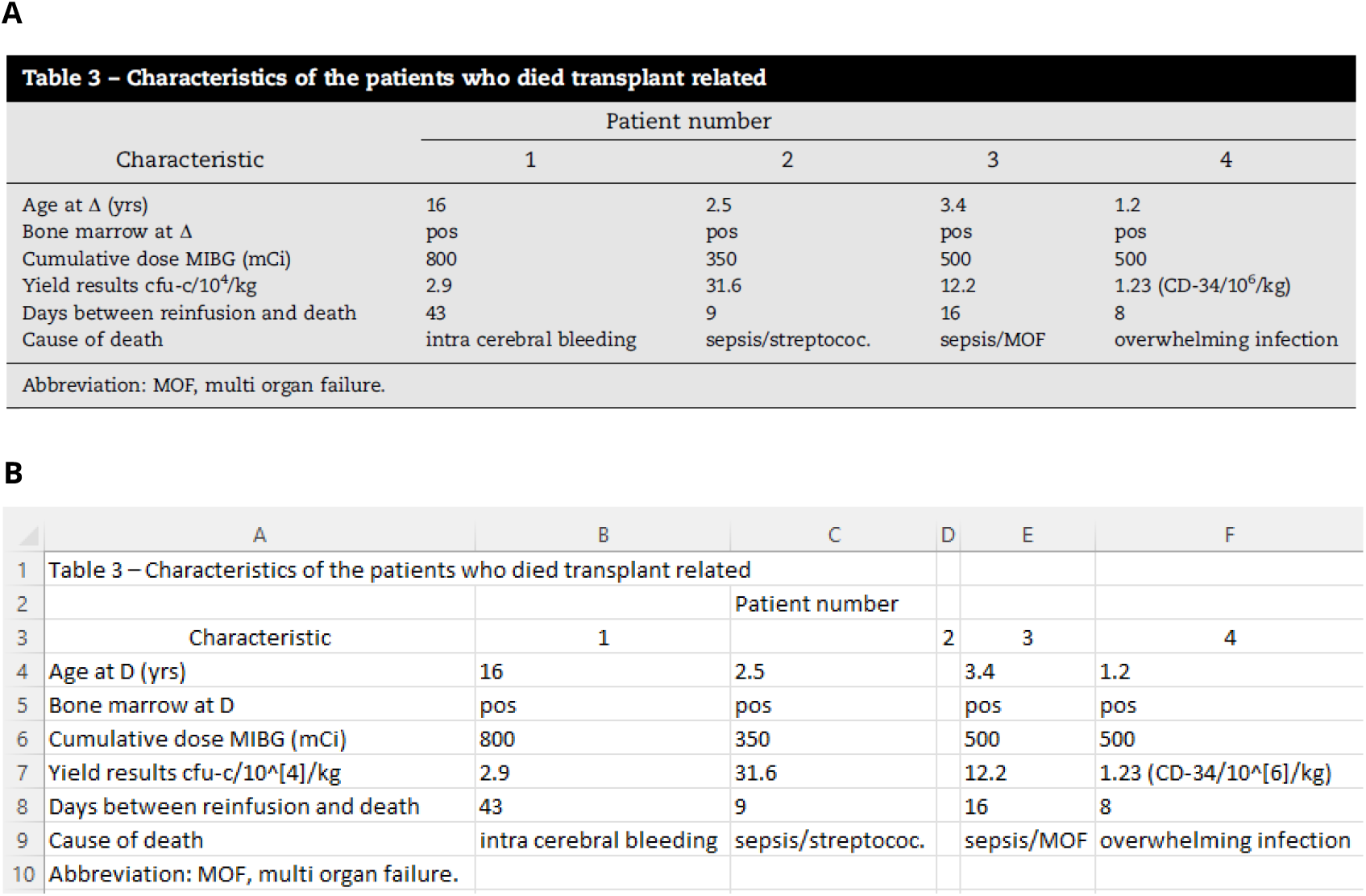
Exemplary results of the pdf2table function included in the PDE_analyzer. The original table 3 (A) was obtained from ***de Kraker et al. (2008*)** and extracted using the PDE_analyzer_i. Merely the column width and text alignment for the PDE-extracted table (B) displayed in Microsoft Excel were adjusted with no modifications to the table content.

## Discussion

A traditional systematic review includes: 1) a generalized literature search in multiple databases with broad search words, 2) an identification of additional records through other sources such as review articles, 3) a manual screening of titles and abstracts after removal of duplicates, and 4) an assessment of full-text articles for eligibility (Fig. 1). Even though there is systematic review software available facilitating the review of titles and abstracts, step three in a traditional approach still is time-intensive, superficial and involves often thousands of articles. For this reason, we developed the PDE R package as pre-screening tool and evaluated its reliability and time saving potential in comparison to the traditional approach used in a Cochrane systematic review on the efficacy and adverse effects of 131I-metaiodobenzylguanidine (131I-MIBG) therapy in patients with newly diagnosed high-risk (HR) neuroblastoma (NBL) by ***Kraal et al. (2017*)**.

Demonstrated through the partial reproduction of a Cochrane Systematic Review (***Kraal et al., 2017*)**, the PDE R package proved to provide tools for quick literature review. With standard parameters, the PDE R package was able to exclude 454/1,263 (35.9%) articles from the required manual review with high sensitivity (Fig. 2) resulting in a significant reduction of review time and reducing the possibility of human errors. Overall, approximately 20% (454/2,291) of the articles identified on PubMed could be excluded by the PDE R package. The PDE R package demonstrated its ability to exclude articles not meeting inclusion criteria. It also sensitively flagged articles for manual review with a large amount of potentially relevant data, in our case clinical data of adverse effects. The package was able to extract tables from PDF files and convert them into an Excel compatible format during pre-screening. Since the PDE R package did not change any PDF files, articles deemed relevant remained compatible with any downstream review softwares such as the Review Manager (RevMan) (***The Cochrane Collaboration, 2020*)** or the EpiTools epidemiological calculator (***Sergeant, 2018*)**.

The PDE R package was specifically designed for user-friendliness and ease of use. To overcome the potential drawback of non-existing graphical user interfaces (GUI) in R (***R Core Team, 2019*)** and the requirement of coding skills, we employed a Tcl/Tk powered GUI. The Tcl/Tk implementation with the tcltk (***R Core Team, 2019*)** and tcltk2 (***Grosjean, 2019*)** packages is part of Microsoft Windows and Linux versions of R by default and easy to install on a Mac as part of XQuartz. Furthermore, the tcltk package offers the display of interactive tables allowing the user to quickly assess the results of the keyword searches. We used the xpdf (***Noonburg, 1996–2020*)** command line tools to allow the searchability of PDF files and the export of relevant tables completely automated. The pdftotext and pdftohtml tools perform especially well in the reassembly of words, e.g., unequally spaced letters or hyphenation at the end of a line.

The PDE R package has the capabilities to enhance the literature search for review articles, gene or disease curation, risk factor analysis, and general literature reviews with its two main functions. The PDE_analyzer function performs the sentence and table extraction on the complete full-text articles. The distinctive features of the PDE_analyzer include 1) filter words, 2) sentence extraction based on search words, 3) abbreviation auto-detection, 4) context extraction, and 5) pdf2table conversion, 6) graphical user interface. The user can provide a list of filter words specific for certain study types or topics. The PDE_analyzer only processes articles which carry words from this list when detected a minimum number of times or at a minimum ration of overall word count as defined by the user (filter word threshold). In addition, filter words and even filter word categories can be used to pre-sort PDF files according to the most common key words. A list of user-defined search words can be used to extract sentences relevant for the evaluation of the suitability of a scientific article. All sentences carrying at least one of the search words are output into a comma- or tab-separated values file, i.e., *.csv or *.tsv. The PDE_analyzer also has the ability to recognize abbreviations of search words used in articles, without the predefinition of any abbreviations being required. The user can choose to export a preferred number of sentences before and after the sentences containing search words and their abbreviations, i.e., select the context. Facilitated by tcltk and tcltk2 R packages, the user can enter analysis parameters using the PDE_analyzer_i interactive version. The interface allows the generation of jobs, execution of analysis and monitoring of active analyses in a visual interface. The user interface does not only enable the rapid selection of parameters and monitoring of active analyses but also the storage of the parameters used for an analysis in a tab-separated values (.tsv) file.

The included PDE_reader permits user-friendly visualization and quick processing of the obtained results. In the additional interface, the user can: 1) quickly browse the sentences with detected keywords, 2) open the full-text article, when required, 3) obtain all tables from the current PDF file in a Microsoft Excel readable file format, 4) and flag or mark articles by adding a prefix, i.e., “!_” or “x_”, to the file name. The prefix can also be easily removed at any time. This is especially useful as it does not require a separate documentation file and allows the interruption of evaluations at any time. For the most part, efficacy of the PDE R package depends on 1) the selection of a filter word threshold and 2) the choice of the filter words. As mentioned before, the filter word threshold is an empirically determined variable. Accordingly, it should be set conservatively low to prevent the introduction of false negatives. However, sensitivity analysis showed that 100% of the articles included in the Cochrane Systematic Review were captured even with a filter word threshold of more than three times the default, i.e., up to 72 (default 20). To select filter words for general pre-screening, the users are required to have prior knowledge of the potential outcomes thus limiting new discoveries. However, high throughput full-text searches possible with the PDE R package are especially useful for the identification of specific search words inside notes or methodologies generally not mentioned in the abstract and have the potential of adding another layer of sensitivity to the literature review. As such, we demonstrated that the full-text article by ***Garaventa et al. (1999*)** was not evaluated by (***Kraal et al., 2017*)** while the PDE R package detected clinical data warranting further full-text studies.

While the processing of full-text PDF files with the package was fast, with 4 second per article on average, the PDE-facilitated pre-screening in contrast to abstract- and title-only reviews required a significant amount of time gaining access and downloading full-text PDF files. We recognize that the PDE R package depends highly on the integrity of the PDF file, cannot process secured or image-only PDF files, and cannot extract tables rotated by 90 degrees. Nevertheless, documentation files are created for secured, non-readable or image-only files or tables. In addition, we are aware that the review of full-text articles can only be applied to a subset of the literature as some articles are not readily available.

We demonstrated that the PDE R package compared to other R packages capable of table extraction such as pdftools (***Ooms, 2019*)** or tabulizer (***Leeper, 2018*)** is able of reliable automatic detection of tables including headers and table legends in scientific articles (Fig. 4). Merely one of the columns was separated into two due to an indented column heading, and Greek letters were not converted correctly. The output tables are saved by the PDE_analyzer in universal comma-separated values (.csv) or tab-separated values (.tsv) files, in contrast to a list of word positions (pdftools). The PDE R package and its table extraction capabilities is completely free of use in contrast to other online tools such as https://smallpdf.com/, https://www.adobe.com/acrobat/online/pdf-to-excel.html, and https://pdftables.com/. In addition, it allows high throughput table extraction and is not limited in input file numbers opposed to online tools such as https://pdftoxls.com/, https://docs.zone/pdf-to-excel, and https://www.pdftoexcel.com/. Accordingly, Specific uses not exclusively include extraction of tables listing functional genomic regions for genomic studies, extraction of clinical study results for epidemiological studies, identification of common abbreviations for drug names for the analyses of drug-related articles, search of specific article types, e.g., case-control studies, retrospective vs. prospective studies, based on filter words for systematic literature reviews.

Features intended to be implemented in the future of the PDE R package include the voluntary exclusion of the reference section for filter and search word detection and the ability to interrupt and continue large dataset analysis even after system crashes. Furthermore, improvements to the PDE reader are planned which include a title and abstract display window, exportable labels and custom comments.

## Conclusion

In conclusion, the PDE R package accelerated the systematic review of literature and simplified the extraction of tables from PDF files. The PDE R package allows the pre-screening and sorting of articles according to filter words, auto-replaces abbreviations, and enables the easy extraction of sentences and tables with user defined search words. It is easily accessible due to a user-friendly graphical user interface and is cross-compatible among Microsoft Windows, Mac, and Linux systems. Therefore, the PDE R package has wide-applicability in various scientific fields using systematic data collection and literature reviews. Given the broad need of easy data extraction from scientific literature and the user-friendly design of the PDE, our application will help scientists at any stage to perform fast and comprehensive literature reviews and reliably obtain data.

## Methods and Materials

### Assembly of the Full-Text Article Library

For partial reproduction of the published Cochrane systematic review by ***Kraal et al. (2017*)**, we used the MeSH headings and text words described by the group in a PubMed search of relevant articles. We downloaded open access papers using the Pubmed Batch Downloader by Bill Greenwald (***Greenwald, 2019*)**, supplemented with a manual download through PubMed with Texas Medical Center (TMC) library access in portable document format (PDF). We obtained articles with restricted access through the TMC Library using the OpenAthens plugin in EndNote. Only articles accessible and available in English were included.

### Literature Evaluation

Using the corresponding filter in PubMed, we identified review articles and processed them separately from primary research articles. For comparison of the PDE R package results with the traditional systematic review approach, we categorized the 1262 full-text articles obtained in PDF format into articles excluded based on title and abstract, full-text articles assessed for eligibility, and articles included by ***Kraal et al. (2017*)**, utilizing the supplemental table named “Characteristics of excluded studies” provided in the Cochrane Review. We did not re-evaluate the articles not-excluded by the PDE R package, accordingly no statement could be made if Kraal *et al.* missed any articles.

### Parameter selection for PDE implementation

To compare the PDE R package with a traditional manual literature review, we translated efficacy and adverse effects described under the “Types of outcome measures” heading in ***Kraal et al. (2017*)** into specific 23 filter/search words (see Supplemental Table S3). For the exemplary literature review, the standard filter word threshold of 20 was chosen. The 1262 PDF files were separated into full-text articles to be assessed for eligibility and excluded articles using the filter word parameter of the PDE_analyzer_i function by automatically copying them into different directories. This is an empirically determined number obtained from the analyses of several thousand articles. Search words for text and table extraction were chosen to be identical to the filter words, since the sentences and tables with the search words provide the most information on the relevance of an article for a systematic review. Two sentences before and after the sentence containing the search word were extracted. This allowed the evaluation of the context as well as focus on the relevant sentence with the PDE_reader_i. Specifically, words like bone marrow or blood pressure had the potential to be abbreviated. Accordingly, any type of detected abbreviation of the words should be counted as an incidence of the word. The standard pixel deviation for indentations in table columns is 20 and was chosen for this representative analysis accordingly. Additional information on table locations was not required. Specifically, 90 degrees-rotated tables not extracted by the PDE_analyzer were selected to be exported as PNG files. All tables were exported as Microsoft Excel-compatible comma-separated values (.csv) files. To evaluate if all files were correctly processed, additional table and text detection documentation files were exported. Intermediate files are of no relevance and were therefore deleted after analysis. The Supplemental Table S3 comprises the TSV file including parameters used for the analysis with a description for their selection. The any columns outside of variable and value columns are generally ignored by the PDE R package. For detailed parameter description please refer to the PDE R package vignette available on CRAN (https://cran.r-project.org/web/packages/PDE/vignettes/PDE.html).

### Computational Details

The PDF data extractor (PDE) R package is available on CRAN (https://CRAN.R-project.org/package=PDE). For our analysis we used PDE version 1.4.3. The R Shiny dashboard web app was published at https://erikstricker.shinyapps.io/PDE_analyzer through shinyapps.io by RStudio. All scripts were executed on R version 4.2.0 (***R Core Team, 2019*)**. The xpdf 4.02 command line tools were obtained from https://www.xpdfreader.com/download.html.

## Additional files

### Supplementary Table S1

#### Detailed list and analytic results of the PDF files processed by the PDE R package

The table lists all 1262 full-text articles processed with the PDE R package together with the analytical results.

### Supplementary Table S2

#### PDE_analyzer output for keyword search in Garaventa *et al.* 1999

The table lists all extracted sentences which included any of the search words listed in the TSV file. The column “search.word.loc_total” indicates in which sentence of the extracted paragraph the search word was found and how many sentences were extracted in total separated by an underscore.

### Supplementary Table S3

#### Parameter file (TSV file) used for the reproduction of Kraal *et al.* (2017) with the PDE R package

The TSV file includes an additional description column guiding the reader through the reasoning for choosing certain parameter values. However, the additional column is ignored by the PDE analyzer and accordingly, the TSV files can be used to recreate the analysis outlined in this article.

## Author Contributions

Conceptualization, E.S. and M.E.S.; methodology, E.S.; software, E.S. and M.E.S.; validation, E.S.; formal analysis, E.S.; investigation, E.S.; resources, M.E.S.; data curation, E.S.; writing—original draft preparation, E.S.; writing—review and editing, E.S. and M.E.S.; visualization, E.S.; supervision, M.E.S.; project administration, M.E.S. All authors have read and agreed to the published version of the manuscript.

## Institutional Review Board Statement

Not applicable.

## Data Availability Statement

Data generated for this publication is available as Supplemental Table S2. The code for the PDE R package is available from the Comprehensive R Archive Network at https://CRAN.R-project.org/package=PDE.

## Acknowledgements

The authors want to thank Tommy H. Tran, Andrea I. Lee, Katherine P. Lee, Erin C. Gregory-Peckham, and Jeremy Schraw for their help in testing the PDE R package, troubleshooting problems and providing valuable feedback for the features.

